# Wolbachia promotes its own uptake by host cells

**DOI:** 10.1101/2022.01.24.477646

**Authors:** Lindsay Nevalainen, Irene L.G. Newton

## Abstract

*Wolbachia pipientis* is an incredibly widespread bacterial symbiont of insects, present in an estimated 25-52% of species worldwide. *Wolbachia* is faithfully maternally transmitted both in a laboratory setting and in the wild. In an established infection, *Wolbachia* is primarily intracellular, residing within host-derived vacuoles that are associated with the endoplasmic reticulum. However, *Wolbachia* also frequently transfer between host species, requiring an extracellular stage to their life cycle. Indeed, *Wolbachia* has been moved between insect species for the precise goal of controlling populations. The use of *Wolbachia* in this application requires we better understand how it initiates and establishes new infections. Here we designed a novel method for live-tracking *Wolbachia* during infection using a combination of stains and microscopy. We show that live *Wolbachia* are taken up by host cells at a much faster rate than dead *Wolbachia* cells, indicating that *Wolbachia* play a role in their own uptake and that *Wolbachia* colonization is not just a passive process. We also show that the host actin cytoskeleton must be intact for this to occur, and that drugs that disrupt the actin cytoskeleton effectively abrogate *Wolbachia* uptake. The development of this live infection assay will assist in future efforts to characterize *Wolbachia* factors used during host infection.

## Introduction

*Wolbachia* are alpha-proteobacteria, part of the anciently intracellular *Anaplasmataceae*, and related to the important human pathogens *Anaplasma, Rickettsia* and *Ehrlichia* (1). However, *Wolbachia* do not infect mammals, but instead are well known for their reproductive manipulations of insect populations, inducing phenotypes such as male-killing, feminization, or sperm-egg incompatibility (2–7). *Wolbachia* is an incredibly successful infection; the symbiont is both vertically and horizontally transmitted and manipulates host reproduction to persist in populations (2, 5, 8–10). In the last decade, *Wolbachia* have been shown to provide a benefit to insects, where the infection can inhibit RNA virus replication within the host, a phenomenon known as pathogen blocking (11). Because insects are vectors for disease, and *Wolbachia* alter the ability of these vectors to harbor important human pathogens, *Wolbachia* are being used to control the spread of diseases such as dengue (12, 13). For *Wolbachia* to establish itself in an insect population it must invade host cells, persist during infection, and be transmitted to the next generation. Therefore, a prerequisite for the use of *Wolbachia* in vector control in insects is the establishment of infection.

Although *Wolbachia* cannot be cultivated *ex vivo*, they can survive for weeks in cell culture media when removed from host cells (2). Additionally, *Wolbachia* can be found in host hemolymph, suggesting that during natural infection of whole organisms, they can be found outside of host cells (2). However, *Wolbachia* entry into the host cell is still relatively uncharacterized. What little we do know about the infection process comes from elegant trans-well assays where *Wolbachia* passively exiting host cells, then passing through a 10 mm barrier, were able to colonize uninfected *Drosophila* cells (14). The infection timeline calculated based on this approach between 3 hours and 6 days, a relatively long time course for most of the *Anaplasmataceae* (15). In that same work, the authors investigated the importance of clathrin-mediated endocytosis and showed that this host process alone cannot account for all *Wolbachia* entry. In this study, we therefore had two goals: to determine *Wolbachia’s* contribution to host uptake and to identify a more precise timeline for direct infection of host cells.

Using both infected and uninfected *Drosophila melanogaster* JW18 cells, we designed a protocol that allows for live visualization of *Wolbachia* establishment. We first asked if live *Wolbachia* are taken up by host cells more rapidly than *Wolbachia* that had been killed. We then established a timeline and quantified the relative numbers of live vs. killed *Wolbachia* entering host cells. Our results suggest that *Wolbachia* indeed do contribute to their own uptake, with live *Wolbachia* colonizing cells more quickly and at higher infection rates than their dead counterparts. We then treated uninfected JW18 cells with cytochalasin-D, which inhibits the polymerization of filamentous actin, and challenged them with live *Wolbachia*. We also show that pre-treatment of host cells with the actin inhibiting drug cytochalasin-D inhibits the uptake of live *Wolbachia* dramatically, identifying the host actin cytoskeleton as important for the internalization process. We therefore conclude that *Wolbachia* are not only passively taken up by *Drosophila* but also actively facilitate uptake.

## Materials and Methods

### Insect Cell Culture

*Drosophila melanogaster* JW18 cells were tetracycline treated to remove *Wolbachia* infection. Infection status was confirmed by PCR after four passages in the presence of antibiotic. Cells with and without *Wolbachia* were maintained in the dark, in T25 unvented cell culture flasks at room temperature. Schneider’s Insect Medium was used and supplemented with 10% heat-treated Fetal Bovine Serum and 1% α/α (penicillin/streptomycin) solution.

### *Wolbachia* Isolation and Overlay for Rate of Uptake Screen

To visualize the rate of uptake of *Wolbachia*, ~2000 uninfected JW18 host cells were seeded into a CellVis 96 well glass bottom plate with fresh Schneider’s media and incubated in the dark at room temperature overnight to allow them to settle. The following day, a confluent T25 flask of JW18 *Wolbachia*+ cells were first counted, then transferred into a 15ml conical tube containing 500μl sterile glass beads and SYTOX Orange (ThermoFisher) dye (250nM). To release *Wolbachia* from JW18 host cells, the conical tubes containing glass beads were then vortexed for 4 pulses of 5 seconds each. Lysate was divided into 500μl aliquots, and each placed onto an Ultrafree MC-SV Durapore PVDF 5.0μm centrifugal filter column (Millipore). Columns were spun at 20,000g at 10°C for 10 minutes to purify *Wolbachia* from host cells (2). Resulting pellets were then transferred into a new 1.5ml centrifuge tube containing fresh Schneider’s media and resuspended. Extractions from the same T25 were divided such that half would be used immediately (live) and the other frozen at −20°C for at least 3 days (killed) to serve as a control for the infection. To prevent bleaching, 50μl of ProLong Live (ThermoFisher) solution was added to each tube (250μL volume total) prior to imaging then incubated away from light for 15min, as per manufacturer’s instructions. JW18 cells are not all infected at the same density (16, 17), and because live *Wolbachia* cannot be quantified by qPCR, we controlled for the multiplicity of infection by counting the number of infected JW18 host cells that we lysed before isolating *Wolbachia*, and by using the matched extracted *Wolbachia* sample as a dead control for each experiment. The *Wolbachia* were then added to each seeded well at an MOI of 20:1 host cell equivalents. That is, for each seeded JW18 cell, we overlayed the *Wolbachia* contents from 20 infected JW18 cells. The plate was then gently spun down for 5min at 500rpm to ensure that *Wolbachia* and uninfected JW18 host cells were proximal. For cytochalasin D treatments, the same infection protocol was followed but preceding overlay, host cells were treated with 15 μg/mL cytochalasin-D in Schneider’s media for 30 minutes.

### Live/Dead Staining of Isolated *Wolbachia*

*Wolbachia* isolated from infected JW18 cells were stained with a 1:1 mixture of SYTO9 (ThermoFisher) and Propidium Iodide (ThermoFisher) either immediately following isolation (live) or after storage at −20°C for at least 3 days prior to imaging (dead). Stained cells were incubated away from light for 20min prior to imaging using the same magnification, intensities, and exposure times.

### Microscopy and Analysis of Images

All images were obtained using a Nikon Ti2 Eclipse inverted microscope at 100x magnification with a Nikon DS-Qi2 camera attachment. Intensities and exposure times are as follows: green/FITC for 200ms with SOLA pad at 4%, red/Texas Red for 400ms with SOLA pad at 8%. All Z stacks were taken with a step size of 0.5μm.

Nikon NiE Elements software was used to first deconvolute all Z stacks (Automatic 3D Deconvolution) and a NiE Elements GA3 program was used to collect internalized *Wolbachia* data using the threshold parameters specified in **Supplemental Table 1** for each image.

Counts and volume data from Nikon NiE Elements were tested for normalcy and overdispersion in R and a zero-inflated GLM with a Poisson distribution (zeroinfl in the PSCL library) was used to test for significant differences in the number and volume of total *Wolbachia* inside of host cells over time in each condition. A logistic regression (*glm* with *family = binomial*) was used to predict infection status of cells based on *Wolbachia* volume and number.

## Results

We were able to successfully isolate live *Wolbachia* from JW18 cells and using a freeze method we were able to kill the population of bacteria for our controls. The efficacy of this method was visualized using live/dead staining (**Fig. 1**). No propidium iodide staining was observed for live, freshly-isolated *Wolbachia* but all cells subjected to freeze-treatment internalized the dye, suggesting the cell membranes were now permeable to propidium iodide staining. Each of the samples were used in live microscopy during infection of *Drosophila melanogaster* cells. Briefly, *Wolbachia* (either live or dead) were stained using SYTOX dye and overlayed onto *Wolbachia-* free JW18 cells. The process was visualized using ProLong Live reagent and fluorescent light microscopy and pictures were taken of live cells every two hours for 4 hours. Live *Wolbachia* could readily be observed entering host cells by 2 hours post infection, whereas we rarely observed internalized killed *Wolbachia* (**Fig. 2**). This result confirmed that we could visualize the infection process and that we could quantify *Wolbachia’s* contribution using a killed control.

**Figure 1.**
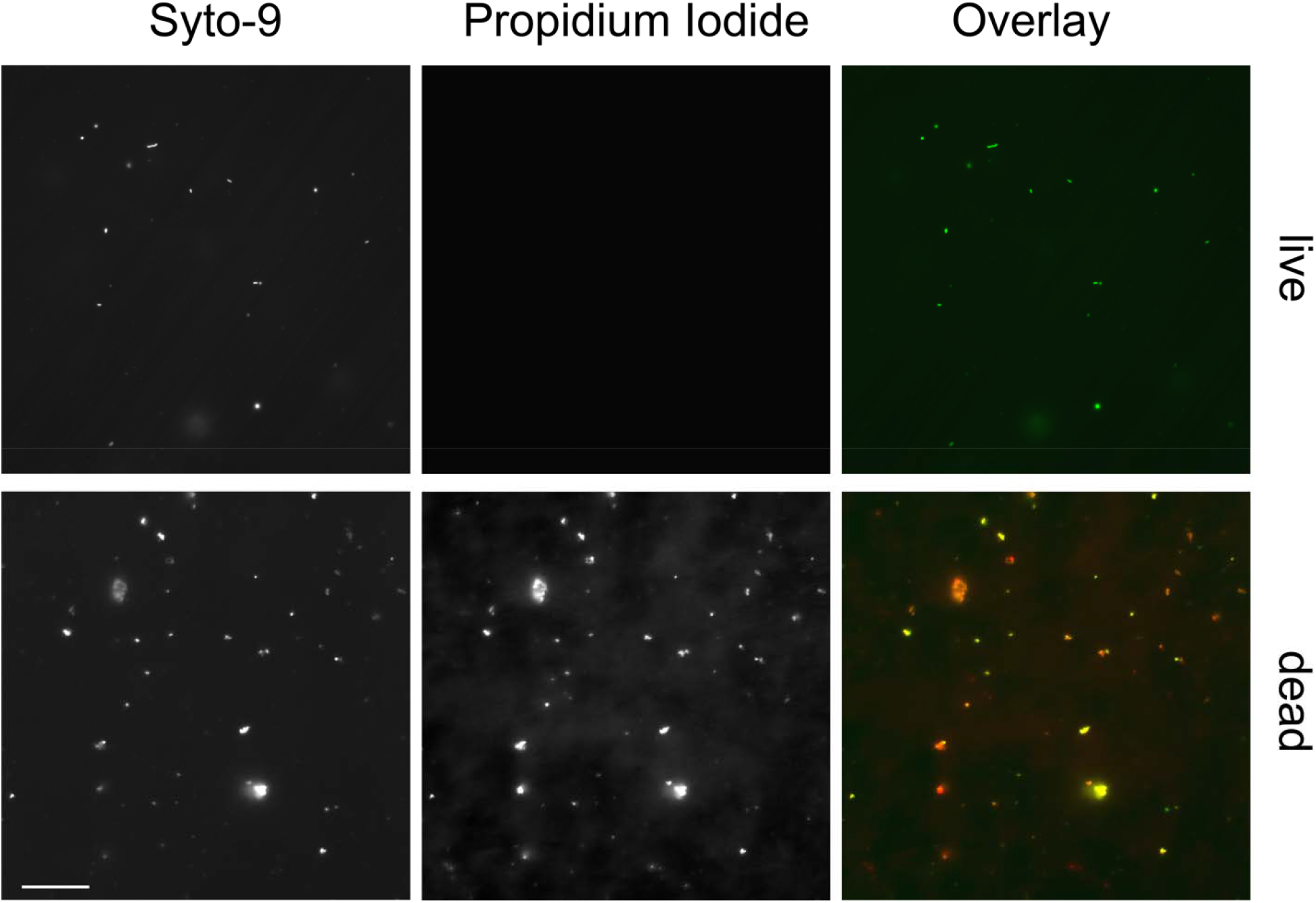
*Wolbachia* are alive after purification from host cells and can be killed with freeze treatment. After isolating *Wolbachia* from *Drosophila* JW18 cells, we stained them using a combination of syto-9 and propidium iodide (PI, see methods). Dead *Wolbachia* are indicated by successful staining using PI. Scale bar = 10 μm.

**Figure 2.**
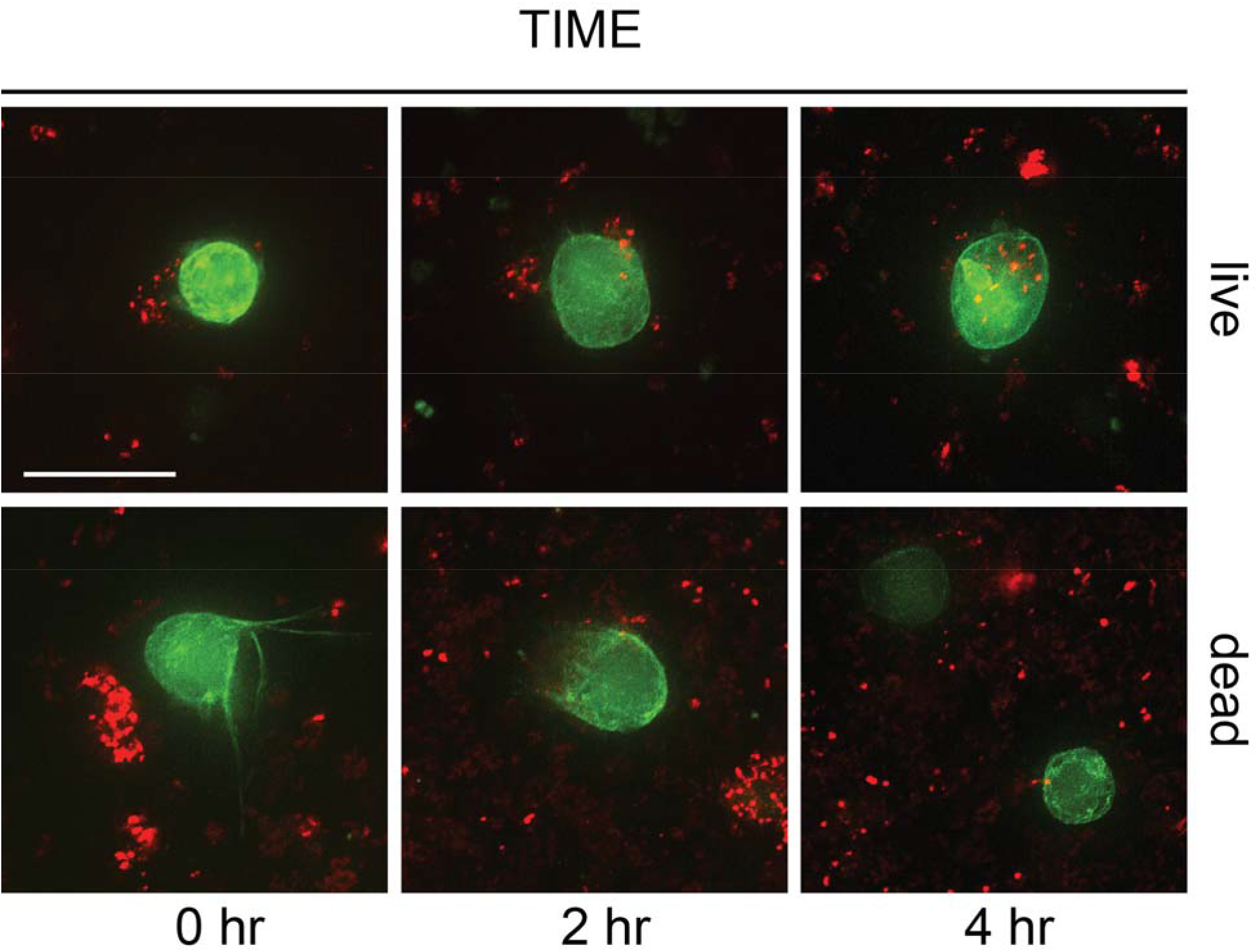
Live *Wolbachia* are readily internalized by *Drosophila melanogaster*. *Wolbachia* (red) stained with SYTOX are visualized infecting JW18 cells (which express Jupiter-GFP, which localizes to microtubules). Representative images during a 4-hour time course are shown. Cell perimeters are delineated by Nikon software using GFP signal intensity across Z-stacks and *Wolbachia* location within the 3-D reconstructed cell identified using SYTOX stain signal. *Wolbachia* were more readily identified within host cells when live compared to killed. Scale bar = 10 μm.

We next proceeded to perform 3 paired experiments, using live and killed counterparts, and counted a total of 454 cells during three four-hour infection time courses (Table 1). Experiments were paired such that the same *Wolbachia* isolated in one experiment were freeze-killed and used as a control for that same experiment. No effect of experiment on the number and volume of *Wolbachia* found within host cells over time was observed (ANOVA; df = 2, F = 1.368, p = 0.256 and df =2, F = 0.311; p = 0.733). However, out of an abundance of caution, and to ensure we could directly compare MOIs, we analyzed the data in aggregate as well as in pairwise fashion, comparing each heat-killed control to each live sample. The two measures we took were of *Wolbachia* counts within cells (individual bacteria-shaped objects) and *Wolbachia* volumes (the calculated volumes, based on pixels, for each of those objects). Both measures were significantly correlated with each other (**Fig. 3A**; Pearson correlation *R = 0*.92; p < 2.2e-16). We therefore present data from the *Wolbachia* counts in the main text (please see **Supplementary Figure 1** for volume-based data).

**Table 1.** Count of host cells, *Wolbachia* volume and mean number for each condition over time.

|  | *Time* | *Host cell number* | *Wolbachia mean volume (s.d.)* | *Wolbachia mean number (s.d.)* |
| --- | --- | --- | --- | --- |
| *Dead* | 0 hr | 54 | 0.03 (0.11) | 0.5 (1.6) |
|  | 2 hr | 77 | 0.24 (1.20) | 2.97 (11.5) |
|  | 4 hr | 78 | 0.10 (0.24) | 2.21 (3.64) |
| *Live* | 0 hr | 81 | 0.25 (1.42) | 3.42 (16.2) |
|  | 2 hr | 96 | 0.80 (2.39) | 13.5 (38.6) |
|  | 4 hr | 68 | 0.52 (0.98) | 8.31 (14.8) |

**Figure 3.**
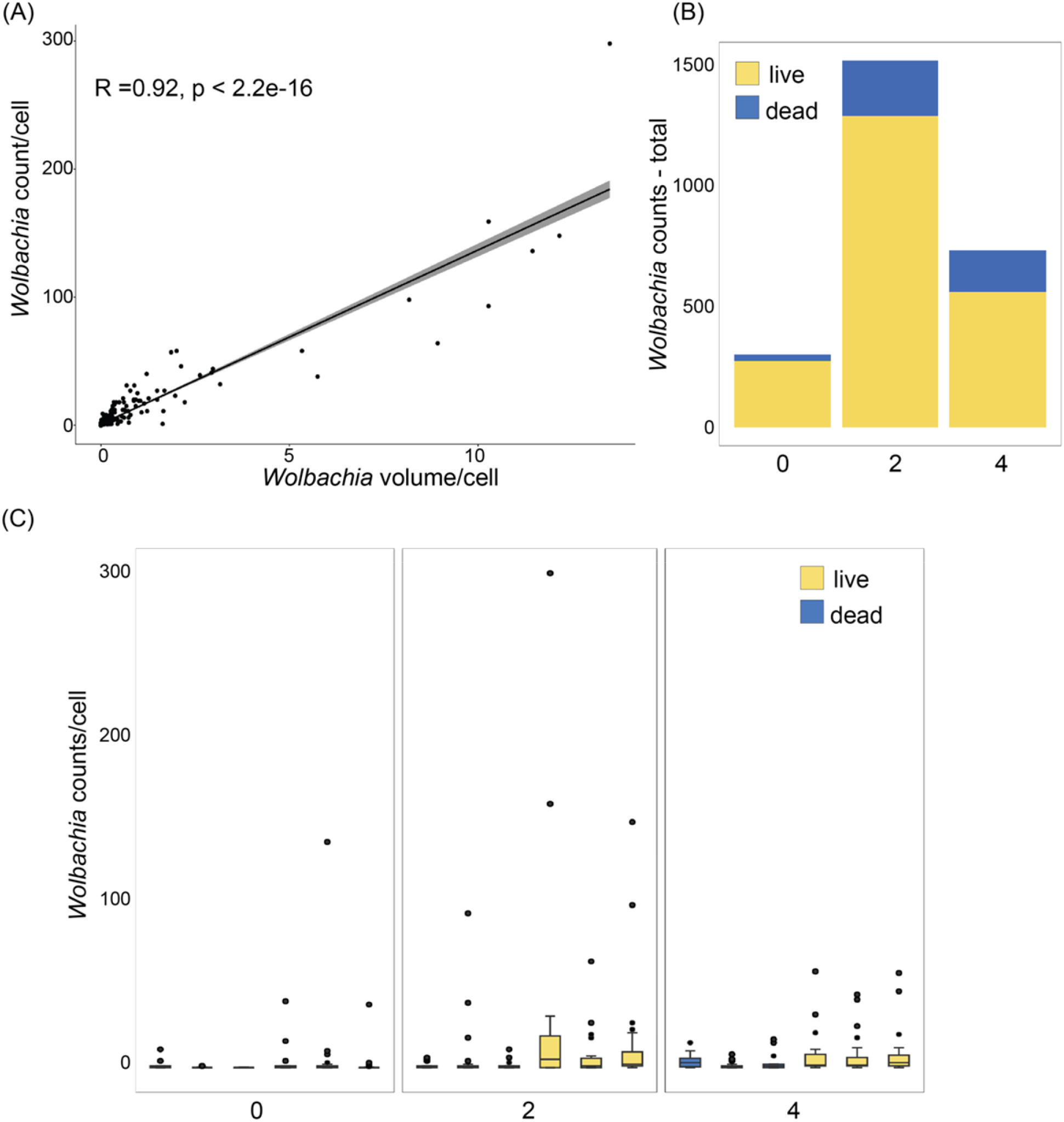
Live *Wolbachia* are internalized more quickly and at higher numbers. Numbers calculated based on live imaging of SYTOX stained *Wolbachia* over a four-hour time-course. (A) Calculated *Wolbachia* numbers and volume per host cell are highly correlated. (B) Aggregate internalized *Wolbachia* counts over the time-course are significantly higher when *Wolbachia* are live vs. dead (χ^2^= 5.6407, df = 1, p = 0.0175). (C) Across three independent experiments, we consistently observed higher *Wolbachia* counts per cell when *Wolbachia* was live relative to heat-killed (dead) matched samples (zero-inflation GLM coefficient estimate −0.4736, z = −2.464, p = 0.137).

### Live Wolbachia are internalized more quickly and at higher numbers than their dead counterparts

Over the course of the infection, we saw an increase in the total number and volume of *Wolbachia* internalized per cell (with median peaks at around 13 and3 in the live vs. dead *Wolbachia* overlays, respectively) (**Table 1**). In aggregate, across all the sampled cells, the number of total *Wolbachia* internalized over the time course of infection was greatest for the treatment with live *Wolbachia* (Fig. 3B; χ^2^= 5.6407, df = 1, p = 0.0175). Indeed, a logistic regression significantly predicts infection status based on both time (estimate = 0.507, stderr = 0.070, z = 7.256, p = 3.89e-13) and whether the assay involved live *Wolbachia* (estimate = 0.723, stderr = 0.211, z = 3.429, p = 0.0006). For two of the three experiments, the largest number and volume of internalized *Wolbachia* was observed in the live overlay after 2 hours (Fig. 3C; 23.1 and 12.4 *Wolbachia*, on average, observed within a host cell). The numbers stabilized at an average of 8.3 *Wolbachia* per host cell at 4 hrs (Table 1).

Because our data are necessarily zero-inflated (due to the large number of uninfected cells in the dead overlay and at early timepoints), we used a zero-inflated GLM to compare number of *Wolbachia* internalized across time for each condition. We observed a significant effect of both time (estimate = −0.079, stderr = 0.014, z = −5.707, p = 1.15e-08) and whether the bacteria were live or dead (estimate = −0.474, stderr = 0.192, z =−2.464, p = 0.0137) on number of *Wolbachia* internalized, corroborating our observations.

### Cytochalasin treatment reduces Wolbachia uptake

Previously, *Wolbachia* has been shown to use the host cytoskeleton for localization within oocytes and for maternal transmission (5, 10). Therefore, we sought to confirm the fraction of uptake of *Wolbachia* we observed that was based on the host cell cytoskeleton. We treated uninfected JW18-tet cells with cytochalasin prior to infection with *Wolbachia* and, as expected, observed a dramatic decrease in internalization (**Fig. 4A**). The median counts observed were generally 1 or zero for each time point (mean of *Wolbachia* within host cells per time; 0hrs μ = 2.33; 2hrs μ = 0.167; 4 hrs μ = 1.62). Indeed, when compared to overlays using dead *Wolbachia*, the cytochalasin D treated samples are statistically indistinguishable (estimate = − 2.055, stderr = 1712.3, z =−0.001, p = 0.950) (**Fig. 4B**).

**Figure 4.**
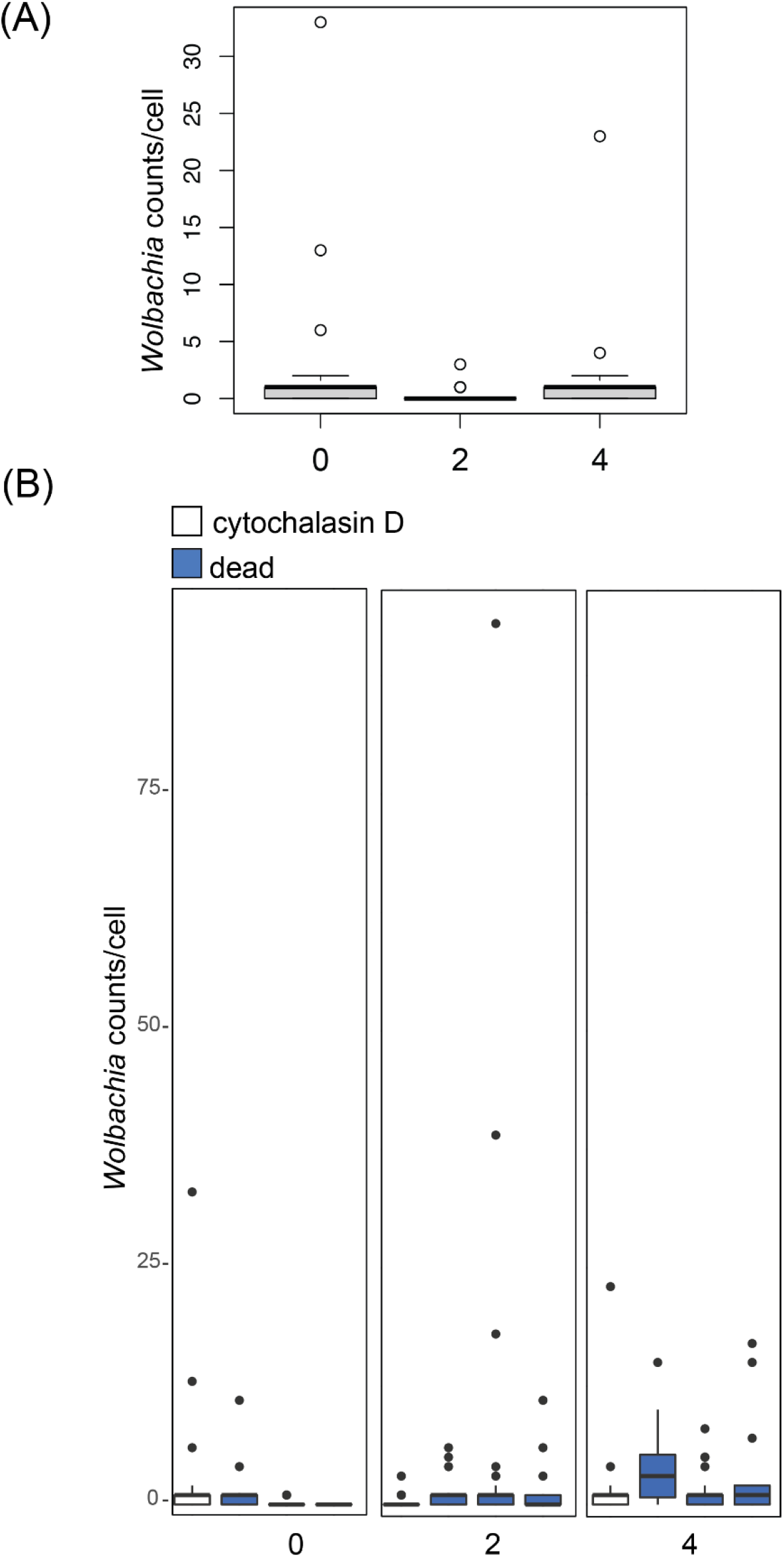
Cytochalasin D treatment reduces the number of internalized *Wolbachia*. Numbers calculated based on live imaging of SYTOX stained *Wolbachia* over a four-hour time-course after treatment of host cells with 15 ug/mL of Cytochalasin D, a toxin that inhibits actin polymerization. (A) *Wolbachia* counts are rarely higher than zero across all timepoints for host cells treated with cytochalasin D and do not vary based on timepoint. (B) The numbers of internalized *Wolbachia* in cytochalasin D treated host cells compared to three independent experiments, using dead *Wolbachia* (zero-inflation GLM coefficient estimate −0.4736, z = −2.464, p = 0.137).

## Discussion

All intracellular bacteria must manipulate the host to enter, persist, replicate, and be transmitted. For those that move between hosts, they must be able to survive extracellularly and reinfect host cells. *Orientia tsutsugamushi*, which causes scrub typhus, utilizes clathrin-mediated endocytosis to enter a host cell, escapes the endosome and utilizes dynein and microtubules to transport itself to the nucleus for replication, then exits the host cell enveloped in host membrane to continue its life cycle (18). For the endosymbiont *Wolbachia*, little is known about its *ex vivo* lifestyle, although more than a decade ago, *Wolbachia pipientis* strain *w*AlbB was observed to survive outside of host cells for one month and be infectious at least 7 days after extraction from the host (19). Additionally, *Wolbachia* can be purified from *Drosophila* host hemolymph and used to inject fruit fly abdomens, after which they establish the canonical infection of the ovaries, suggesting that the microbe has a free-living form conducive to horizontal transfer (20). Understanding how *Wolbachia* moves between hosts and establishes infection is a primary research goal for the use of *Wolbachia* in vector control.

Here we used cultured *Drosophila* cells to identify a timeline of *Wolbachia* entry into host cells, and to characterize *Wolbachia’s* role in its own uptake. We show that a population of live *Wolbachia* can directly infect established cultured JW18 host cells in as few as 2 hours (**Fig. 2, Fig. 3**). We show that this population of live *Wolbachia* enters host cells much more rapidly, and in greater numbers compared to killed controls (**Fig. 2, Fig. 3, Table 1**). This strongly implies that *Wolbachia* play a role in their own uptake; if this was purely a passive process, the live and dead populations would enter host cells at comparable rates and in similar numbers. This work confirms a previous study that has shown that heat-killed *Wolbachia* directly placed into uninfected cell culture are not detectable by either fluorescence in-situ hybridization (FISH) or PCR after incubation for 5 days, while live *Wolbachia* are easily detectable by both methods after this period of time (19). Our results are the first to show a direct comparison between the rates of initial colonization of live and dead *Wolbachia* by direct infection in cell culture in less than 24 hours.

Previous work has established that *Wolbachia* interacts with the host actin cytoskeleton, both requiring it for efficient host colonization and transmission and using a secreted effector that bundles host actin to facilitate transmission (10, 21). Work from the Sullivan lab clearly established that *Wolbachia* utilize dynamin and clathrin-mediated endocytosis to enter host cells, but inhibition of these pathways did not prevent entry completely (22). This indicates that *Wolbachia* may utilize redundant routes of entry. To build on and expand the known *Wolbachia* routes of entry, we next endeavored to show that the host cell actin cytoskeleton is needed for this initial entry process to occur. We treated uninfected JW18 cells with the drug cytochalasin-D, then exposed these treated cells to live *Wolbachia* as before. Importantly, live *Wolbachia* were unable to enter the treated cells (**Fig. 4**), and indeed we observed numbers even more reduced compared to the dead *Wolbachia* controls. This result suggests that the host actin cytoskeleton must be functional for the initial colonization step to occur.

When *Wolbachia* finds itself outside of host cells, it may alter its protein expression, likely to allow it to survive extracellularly. Understanding the changes in *Wolbachia* gene expression during *ex vivo* growth and during host infection will help identify *Wolbachia* loci that are important for host internalization. Many bacterial pathogens, for example *Coxiella*, use acidification of the vacuole to detect host uptake, and it may be that *Wolbachia* similarly responds to the acidification of its host-derived vacuole after uptake.

In conclusion, this work shows that *Wolbachia* promotes its own uptake by host cells, which helps elucidate how this bacterial symbiont has become such a globally successful infection. We also show that the host actin cytoskeleton is necessary for *Wolbachia* uptake, which further expands knowledge necessary to use the microbe in vector control efforts.

## Acknowledgements

We thank Dr. Bill Sullivan for the JW18 cell lines used in this work and Newton laboratory members for their feedback on early drafts of this manuscript. This work was supported by NIH grant R01 AI144430 to ILGN.

---

**Supplementary Figure 1. Live** *Wolbachia* **are internalized more quickly when volume is used to quantify***Wolbachia* **load.** Numbers calculated based on live imaging of SYTOX stained *Wolbachia* over a four-hour time-course. Across three independent experiments, we consistently observed higher *Wolbachia* volumes per cell when *Wolbachia* was live relative to heat-killed (dead) matched samples.

